# A lamin A/C variant causing striated muscle disease provides insights into filament organization

**DOI:** 10.1101/2020.10.20.347088

**Authors:** Rafael Kronenberg-Tenga, Meltem Tatli, Matthias Eibauer, Wei Wu, Ji-Yeon Shin, Gisele Bonne, Howard J. Worman, Ohad Medalia

**Affiliations:** Department of Biochemistry, University of Zurich, Winterthurerstrasse 190, 8057 Zurich, Switzerland; Department of Medicine and Department of Pathology and Cell Biology, Vagelos College of Physicians and Surgeons, Columbia University, New York, NY; Sorbonne Université, INSERM, Centre de Recherche en Myologie, Institut de Myologie, F-75651 Paris CEDEX 13, France

## Abstract

The *LMNA* gene encodes the A-type lamins that polymerize into ~3.5 nm thick filaments, and together with B-type lamins and lamin binding proteins form the nuclear lamina. Mutations in *LMNA* are associated with a wide variety of pathologies. In this study, we analyzed the nuclear lamina of embryonic fibroblasts from *Lmna*^H222P/H222P^ mice, which develop cardiomyopathy and muscular dystrophy. Although the organization of the lamina appeared unaltered, there were changes in chromatin and B-type lamin expression. An increase in nuclear size and consequently a relative reduction in heterochromatin near the lamina allowed for a higher resolution structural analysis of lamin filaments using cryo-electron tomography. This was most apparent when visualizing lamin filaments *in situ*, and using a nuclear extraction protocol. Averaging of individual segments of filaments in *Lmna*^H222P/H222P^ mouse fibroblasts resolved two-polymers that constitute the mature filaments. Our findings provide better views of the organization of lamin filaments and the effect of a striated muscle disease-causing mutation on nuclear structure.

## Introduction

Nuclear lamins are the intermediate filament (IF) building blocks of the nuclear lamina on the nucleoplasmic aspect of the inner nuclear membrane of metazoan cells (Fisher et al., 1986; McKeon et al., 1986). They are classified as type V IF proteins based on their sequences (Steinert and Roop, 1988). Similar to cytoplasmic IF proteins, lamins contain a long rod domain comprised of four coiled-coil α-helical segments, termed 1A, 1B, 2A and 2B, separated by flexible linkers (Gruenbaum and Foisner, 2015). This domain is flanked by a non-helical N-terminal head and C-terminal tail domain. The C-terminal domain hosts a nuclear localization sequence, and an immunoglobulin (Ig) like fold.

The nuclear lamina primarily provides mechanical support to the cell nucleus (Maurer and Lammerding, 2019; Pfeifer et al., 2019; Sapra et al., 2019). Four main lamin isoforms are found in mammals. In humans, they are encoded by the *LMNA*, *LMNB1*, *LMNB2* genes that in somatic cells encode lamin A/C, lamin B1 and lamin B2, respectively (Worman, 2012). While the B-type lamins are expressed in almost all mammalian cell types, the expression of A-type lamins is developmentally regulated and occurs primarily in differentiated cells (Constantinescu et al., 2006; Kim et al., 2011). Both types of lamins are localized to the nuclear periphery; however, small amounts of A-type lamins are also found in the nucleoplasm where they may function in chromatin organization and gene regulation (Naetar et al., 2017).

In solution, lamin dimers form a parallel coiled-coil structure between two monomers (Klapper et al., 1997). These further assemble by head-to-tail association into long polymers, which further associate laterally into the mature filaments (de Leeuw et al., 2018; Stuurman et al., 1998). However, due to the length and flexibility of the coiled-coil domain, which is ~50 nm, structural determination of lamin dimers has been a challenging task (Makarov et al., 2019; Zwerger and Medalia, 2013). A detailed atomic model of full-length lamin proteins is still elusive. However, structures of lamin A fragments (Ahn et al., 2019; Herrmann and Aebi, 2004; Kapinos et al., 2011; Lilina et al., 2019) have exemplified the interactions between lamin coil 1B to form a tetrameric filament (Ahn et al., 2019; Lilina et al., 2019). The atomic structure of the Ig-fold of human lamin A/C has also been determined, exhibiting a globular two β-sheets structure (Dhe-Paganon et al., 2002; Krimm et al., 2002). The analysis of coiled-coil fragments provided insights into the intra-organization of lamin dimers (Strelkov et al., 2004). Based on such analysis (Ahn et al., 2019) an alternative model for lamin assembly in which lateral interactions between the coiled-coil of the dimers governs the lateral assembly, thus, head-to-tail polymer is avoided. An analysis of lamin filaments in fibroblasts has shown that lamins assemble into 3.5 nm thick filaments within a ~14 nm thick meshwork attached to the inner nuclear membrane (Turgay et al., 2017). The filaments exhibit a short persistence length of <200 nm, which not only hints at their unique mechanical properties and flexibility (Sapra et al., 2019), but also impose a major challenge for structurally reconstruction of lamin filaments.

Mutations in the genes encoding lamins, particularly *LMNA*, cause human diseases termed laminopathies (Tatli and Medalia, 2018; Worman and Bonne, 2007). The most common of these rare laminopathies is dilated cardiomyopathy usually associated with muscular dystrophy (Maggi et al., 2016) (Nicolas et al., 2019). The *LMNA* p.His222Pro (hereafter H222P) missense mutation was first identified in a family with autosomal dominant Emery-Dreifuss muscular dystrophy (Bonne et al., 2000). Homozygous *Lmna*^H222P/H222P^ mice develop dilated cardiomyopathy and regional skeletal muscular dystrophy that phenocopies the human disease (Arimura et al., 2005; Muchir et al., 2012). Cells from *Lmna*^H222P/H222P^ mice have altered stiffness (Chatzifrangkeskou et al., 2020) and abnormalities in several cell signaling pathways (Choi et al., 2018). However, it is not clear how this mutation affects the structure of the nuclear lamina and lamin filaments. The H222P amino acid substitution is localized to the linker domain of lamin A/C, between coil 1B and 2A of the proteins. While substitution with a proline would presumably disrupt α-helix structure, it does not play a role in the predicted coiled-coil interactions.

To obtain insights into how the H222P amino acid substitution affects lamin structure, we used several modalities to analyze fibroblasts from *Lmna*^H222P/H222P^ mice. Quantitative immunofluorescence and total internal reflection fluorescence (TIRF) microscopy indicated nuclear alterations in these cells, while cryo-electron tomography (cryo-ET) provided new structural insights into the nuclear lamina and the lamin filaments *in situ* and in ghost nuclei. We show that the nuclear area is increased while heterochromatin is proportionally reduced. This results in an increase of contrast for lamin filaments using cryo-ET. Applying averaging approaches to *in silico* fragmented filaments (Martins et al., 2020) produced an unprecedented view of the lamin filaments and their substructures. Moreover, mapping back the structural class averages into their original position on lamin filaments provides better resolved insight into lamin filament organization. While the structure of lamins is presumably unaffected by the pathogenic H222P amino acid substitution in lamin A/C, the chromatin organization at the nuclear lamina and overall nuclear organization is altered.

## Results

### The nucleus, lamins and condensed chromatin in *Lmna^H222P/H222P^* fibroblasts

At a cellular level, the impact of amino acid substitutions on A-type lamins on nuclear organization has been previously demonstrated for several point mutations (Bertrand et al., 2020; Goldman et al., 2004; Scaffidi and Misteli, 2008; Vigouroux et al., 2001). To gain insights into the impact of the lamin A/C H222P amino acid substitution on the nucleus, and in particular on nuclear lamin filaments, we examined immortalized *Lmna*^H222P/H222P^ and wild type mouse embryonic fibroblasts (MEFs). Using TIRF microscopic analysis, we confirmed that both lamin A/C and lamin B1 were properly localized to the nuclear lamina in *Lmna*^H222P/H222P^ MEFs similar to wild type, but the nuclei appeared larger (Fig. 1A). Quantitative analysis showed that the nuclei area of *Lmna*^H222P/H222P^ MEFs was significantly increased at 1.09 times that of wild type (Fig. 1B). We used immunofluorescence microscopy to obtain signals to quantify the amounts of histone H3K9me3, a marker of heterochromatin, histone H327ac, a maker of transcriptional activity, lamin B1 and lamin A/C in wild type and *Lmna*^H222P/H222P^ MEFs (Fig. S1). This analysis showed that the *Lmna*^H222P/H222P^ MEFs had significantly decreased histone H3K9me3, increased histone H3K27ac, increased lamin B1 and decreased lamin A/C (Fig. 1B). When analyzed by immunoblotting, lamin B1 and lamin B2 expression was increased in the *Lmna*^H222P/H222P^ MEFs; however, the histone H3K9me3 signal was slightly increased and the lamin A/C signal was not reduced (Fig. S2). These results suggest that incorporation of lamins into the nuclear lamina is not directly affected by the lamin A/C H222P amino acid substitution in linker 1-2, between coil 1 and 2. The increase in nuclear surface area likely resulted in reduction of heterochromatin levels per nuclear envelope area, rather than changes in the absolute quantity of heterochromatin. The changes in lamin expression are presumably attributed to the increased surface of the nuclear envelope that may induce an upregulation of lamin B1, while the overall chromatin and lamin A/C levels resemble their original amounts.

**Fig. 1.**
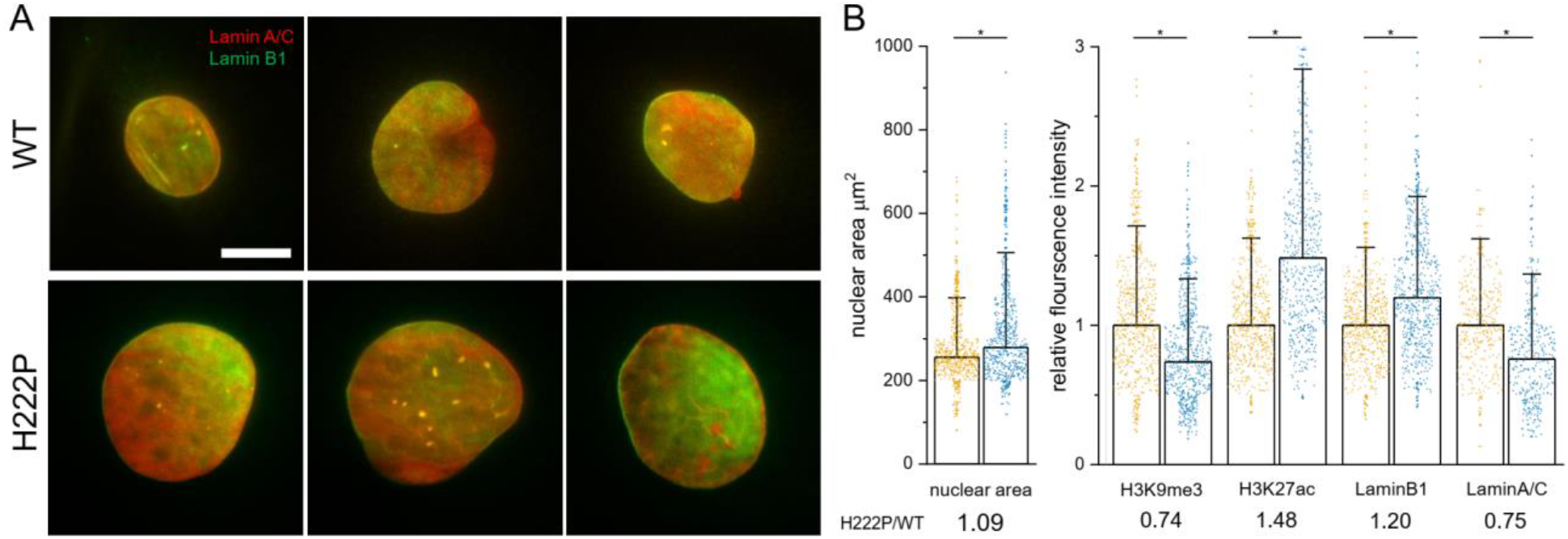
Nuclear organization of*Lmna*H222P/ H222P MEFs. **A.**TIRF microscopy images indicate the localization of lamin A/C and B1 in wild type and *Lmna*^H222P/H222P^ MEFs. **B.** Nuclear area and immunofluorescence microscopy-based quantifications of histone H3K9me3, histone H327ac, lamin B1 and lamin A/C in wild type and *Lmna*^H222P/H222P^ MEFs. Each dot represents the average signal intensity of a single nucleus. The height of the bars is set to the median and the error bar indicates 1.5 standard deviations. Scale bar 10 μm. Number of nuclei analyzed: Nuclear area n=602, H3K9me3 n=602, H3K27ac n=552, lamin B1 n=602, lamin A/C n=356. * p<0.05 between *Lmna*^H222P/H222P^ and wild type by one way ANOVA Tukey’s multiple comparison test.

### Nuclear lamina structure in *Lmna^H222P/H222P^* MEFs

The molecular organization of the nuclear envelope can be visualized by cryo-ET (Harapin et al., 2015; Weber et al., 2019). We cultured cells on an EM-grid prior to vitrification and focus ion beam (FIB) milling in conjunction with cryo-ET (Rigort et al., 2012). This allowed us to visualize the organization of the nuclear envelope in *Lmna*^H222P/H222P^ MEFs *in situ*. The nuclear membranes, lamin filaments and chromatin were all seen without any apparent structural alterations; however, the lamins were better resolved in the mutant cells (Fig. 2A,B; Fig. S3). Surface rendering of the thin reconstructed sections provides a view of the nuclear envelope regions, including the adjacent cytoplasmic structures (Fig. 2C). The nuclear areas hosting lamin filament meshworks were clearly visible with less chromatin and other nuclear components shadowing the lamins in *Lmna*^H222P/H222P^ MEFs compared to wild type ones (Fig. 2C, Fig. S4A,B). The reduced density of nuclear structures at the lamina detected in cryo-tomograms of *Lmna*^H222P/H222P^ nuclear envelopes confirmed the fluorescent microscopy analysis and indicated a bigger gap and reduced interaction of lamins with dense chromatin.

**Fig. 2.**
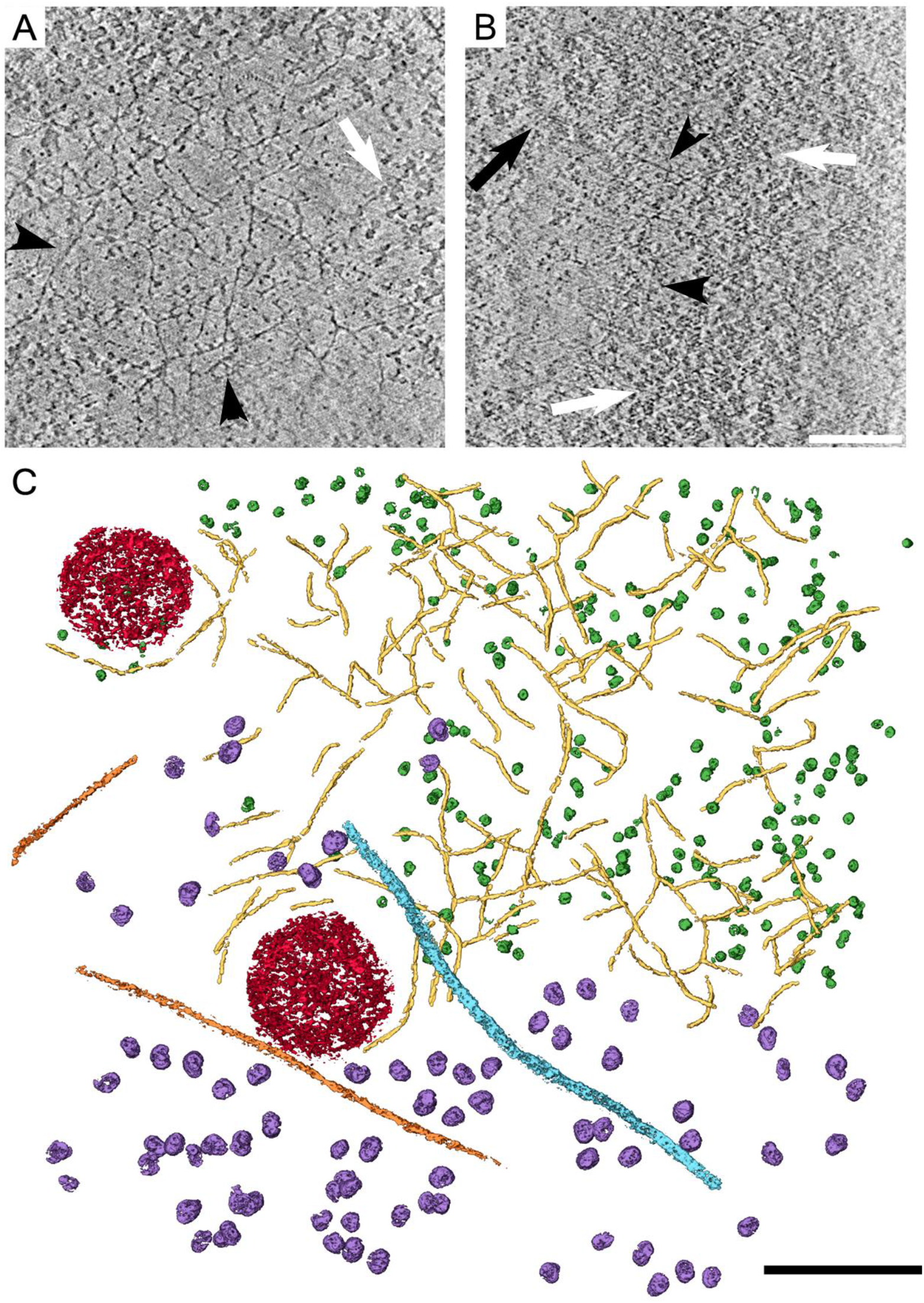
Visualizing the nuclear lamina in *Lmna*_H222P/H222P_ MEFs, *in situ*. **A, B.** Lamin filaments (arrowheads) in an x-y section, 7 nm in thickness, through a tomogram of *Lmna*^H222P/H222P^ (**A**) and wild type (**B**) MEFs. Lamin filaments (arrowheads) are better resolved in the *Lmna*^H222P/H222P^ MEFs. Fewer nucleosomes (white arrows) are in close proximity to the lamina (see Fig. S3). Vitrified cells were subjected to cryo-FIB prior to cryo-ET (see Materials and methods). Black arrow indicates a nuclear pore complex. Scale bar 100 nm **C.** A surface rendering view of a *Lmna*^H222P/H222P^ MEF lamelea, 150 nm in thickness, shows the organization of lamin filaments (yellow), nucleosomes (green), nuclear pore complexes (red), actin (orange), cytoplasmic vimentin filaments (turquoise) and ribosomes (purple). Scale bar 100nm

To further enhance visualization of the nuclear lamins, we knocked down vimentin expression and removed the cytoplasm, using short exposure to mild detergent followed by nuclease treatment prior to rapid vitrification (Turgay and Medalia, 2017). This produced ghost nuclei in which lamin filaments could be readily identified and followed over longer distances and large data sets acquired from most positions along the nuclear envelope (Tenga and Medalia, 2020). Here, we applied this procedure and detected lamin filaments in both *Lmna*^H222P/H222P^ and wild type ghost nuclei (Fig. 3A, B, respectively). The lamin meshworks were better identified in the ghost nuclei from the mutant cells (Fig. 3A). A surface rendering of *Lmna*^H222P/H222P^ MEFs showed lamin filaments, nuclear pore complexes and cytoplasmic vimentin filaments (Fig. 3C) that were similar to that previously reported in wild type fibroblasts (Turgay et al., 2017). The overall organization of lamin filaments in ghost nuclei from *Lmna*^H222P/H222P^ MEFs resembled the wild type nuclear lamina as seen by the rendered view of two tomograms (Fig. S4C,D). While nuclease treatment only removed part of the heterochromatin from ghost nuclei of wild type MEFs, these densities were hardly detected in the ghost nuclei prepared from the *Lmna*^H222P/H222P^ MEFs (Fig. S4C,D, green). These experiments and cryo-ET analysis suggest that the overall organization of the nuclear lamins is not substantially altered in *Lmna*^H222P/H222P^ MEFs. Rather, the H222P amino acid substitution in lamin A/C influences the size of the nucleus and the chromatin organization at the interface of the lamina, providing a better view of the nuclear lamina.

**Fig. 3.**
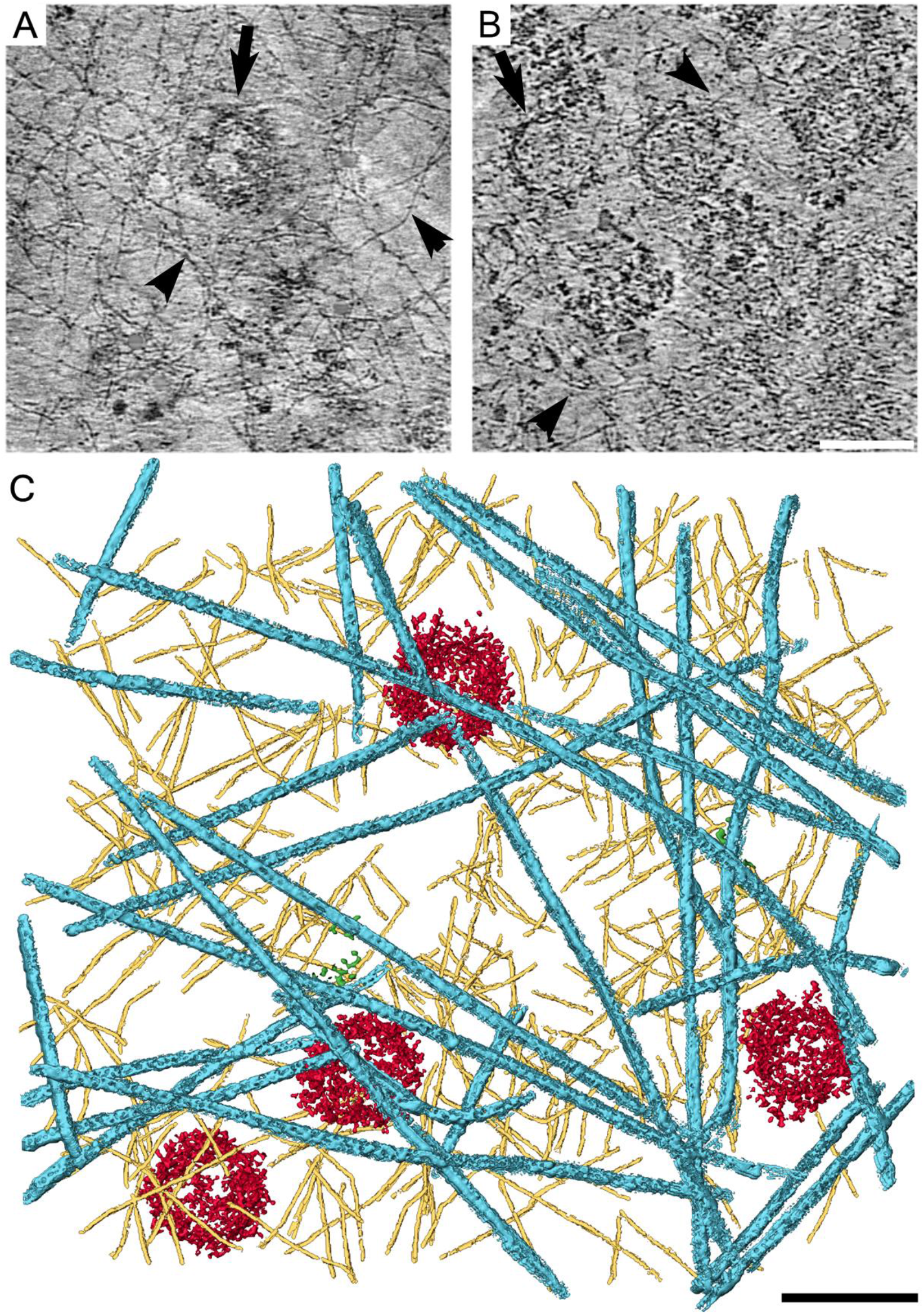
Visualizing the nuclear lamina in *Lmna*_H222P/H222P_ ghost nuclei resolved the nuclear lamina meshwork. **A, B.** Lamin filaments (arrowheads) in an x-y section, 7 nm in thickness, through tomograms of *Lmna*^H222P/H222P^ (**A**) and wild type (**B**) MEFs. Vitrified cells were subjected to cryo-FIB prior to cryo-ET (see Materials and methods). Black arrows indicate nuclear pore complexes. Scale bar 100 nm. **C.** A surface rendering view of *Lmna*^H222P/H222P^ MEFs ghost nuclei preparation, shows the organization of lamin filaments (yellow), nuclear pore complexes (red) and cytoplasmic vimentin filaments (turquoise). Scale bar 100nm

### Structural organization of lamin filaments by averaging and filament reconstitutions

Lamins dimerize in solutions and form head-to-tail polymers (Stuurman et al., 1998); however, how these long coiled-coils protein structures are further assembled into the filaments of the nuclear lamina remains unknown. The higher contrast of individual lamin filaments observed by cryo-ET of *Lmna*^H222P/H222P^ MEFs encouraged us to acquire 250 cryo-tomograms of ghost nuclei to gain further insights into their organization and formation. Lamin filaments in these nuclei appeared flexible and often curved, although some filaments exhibit a straight appearance, and were occasionally decorated with globular densities (Fig. 4A). These filaments closely resembled lamin filaments extracted from wild type MEF ghost nuclei (Fig. S5A). The globular densities were presumably the Ig-like fold domains (Turgay et al., 2017). The position of these globular structures along the filament was roughly regular with some variable distance from the axis of the filaments. This was likely due to the 70 amino acid linker between the end of helix 2B and the Ig-like fold domain. The heterogeneity in the position of Ig-fold domains and heterogeneity of lamin filaments, as well as and the possibility of other proteins binding to the lamins, prevents using conventional averaging approaches to analyze their structure. In order to reduce the complexity and flexibility of these filaments, we therefore fragmented them into short segments (12 nm in length and 4.4 nm in width) *in silico* and calculated their 2D projections followed by a single particle classification approach (Martins et al., 2020). The restricted width of the analyzed areas allowed us to primarily focus on the coiled-coil rod domains that constructed the core of the filaments, with the risk of partial exclusion of the Ig-like fold domains. Analysis of the prominent 2D classes representing different views revealed substructures within the ~3.5 nm diameter lamin filaments, most pronounced among them two ~1.8 nm thick filamentous substructures often interacting and crossing each other and sometimes merging into one structure (Fig. 4B asterisks, Fig. S5B). We next mapped back the class averaged structures of lamin filaments to their original coordinates in the ghost nuclei. This produced a set of reconstituted lamin filaments resolved to higher resolution and with higher contrast (Fig. 4C). Two protofilaments compose the mature filaments interacting and crossing each other frequently but not in a uniform manner (Fig. 4C, asterisks). These substructures resembled in shape and dimensions the head-to-tail polymer of dimers. The presence of two polymer structures within lamin filaments and their heterogenous appearance may increase the spectrum of conformations that can be adopted by the mature filaments. This emphasizes the possible variations within the structure. The interactions between the two head-to-tail polymers may vary to allow tight interactions as well as more scarce appearance, providing additional capability for lamin filaments to fulfill various functions. Resolving the 3D structure of the lamin filaments, together with their Ig-like fold domains, could provide additional information on lamin assembly and the differences between A- and B-type lamin filaments. However, a larger data set would be needed in addition to an innovative image processing approach.

**Fig. 4.**
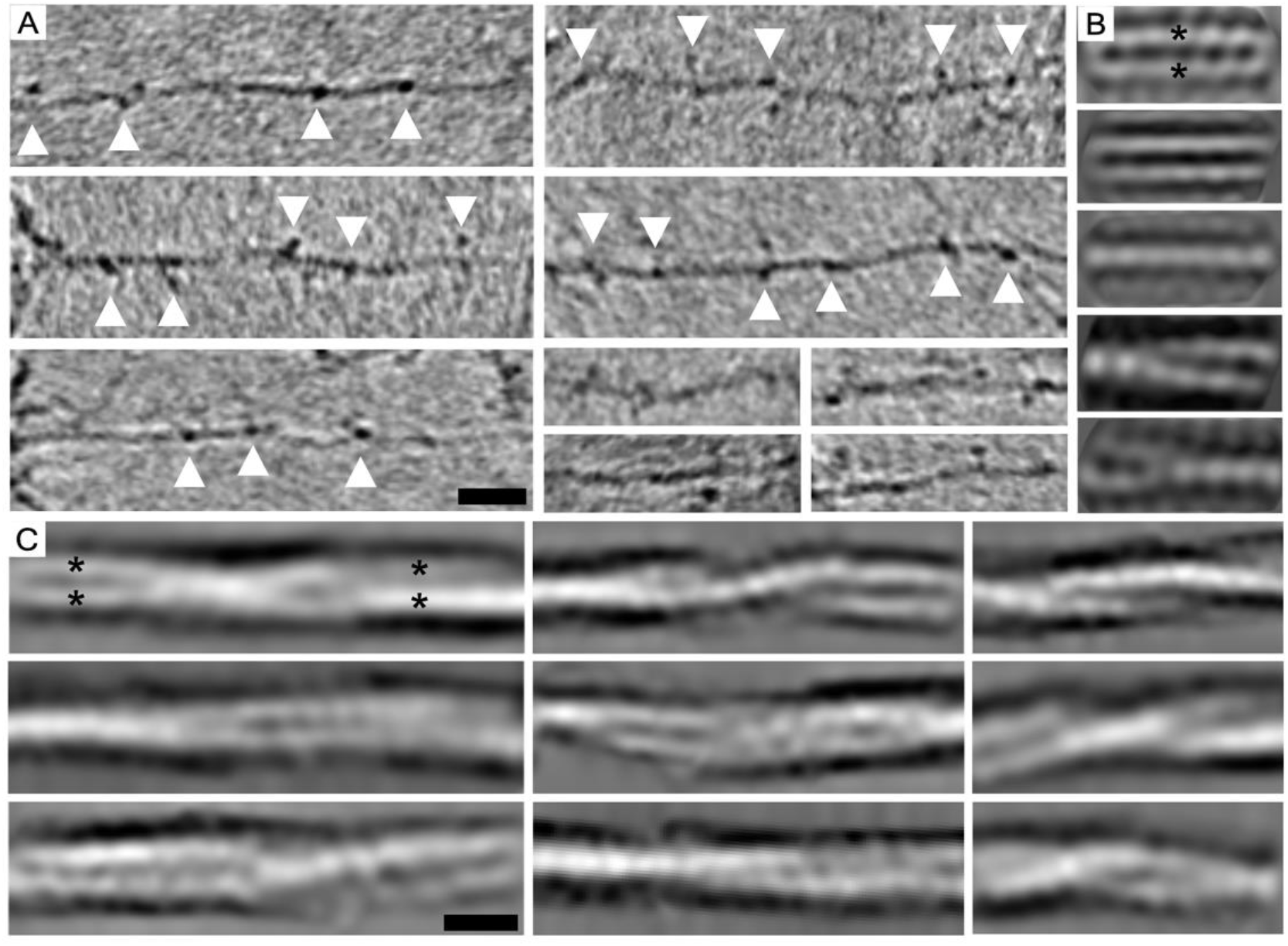
Structural analysis of lamin filaments in ghost nuclei from *Lmna*_H222P/H222P_ MEFs. **A.** A set of individual filaments at the nuclear envelope of from a cryo-tomogram of a ghost nucleus. The filaments are mostly curved and structurally heterogeneous. Globular structures along the lamin filaments (arrowheads) and presumably the Ig-like domains, which exhibit variability in position. Scale bar 20 nm. **B.** Individual two-dimensional class averages, 12 nm long, show different views into lamin filament organization and orientations. All class averages are shown in Fig. S5B. Asterisks indicates two filamentous substructures interacting and crossing each other and even situated one behind the other (middle). **C.** Reconstitute filaments show detailed organization of the filaments and their substructures. The 2D averaged classes (**B**) were used to map-back the averaged structures to form reconstitute filaments with the correct order. Asterisks indicate protofilaments interacting and crossing each other. Scale bar 4 nm.

## Discussion

Based on our novel data, we have generated a model of the nuclear envelope and lamin organization in wild type and *Lmna*^H222P/H222P^ MEFs (Fig. 5). Nuclei of *Lmna*^H222P/H222P^ MEFs have a larger 2D surface area and less densely packed heterochromatin and likely chromatin-associated factors. While the lamin filaments of the lamina of *Lmna*^H222P/H222P^ MEFs are structurally similar to wild type lamin filaments, they are better visualized by cryo-ET, presumably because of decreased interactions with the densely-packed chromatin at the nuclear periphery. The contrast of the lamin filaments in nuclei of *Lmna*^H222P/H222P^ MEFs is substantially higher than the filaments previously observed in wild type vimentin-null MEFs (Turgay et al., 2017). This provides us with a unique system to obtain a better resolved view of lamin filaments.

**Fig. 5.**
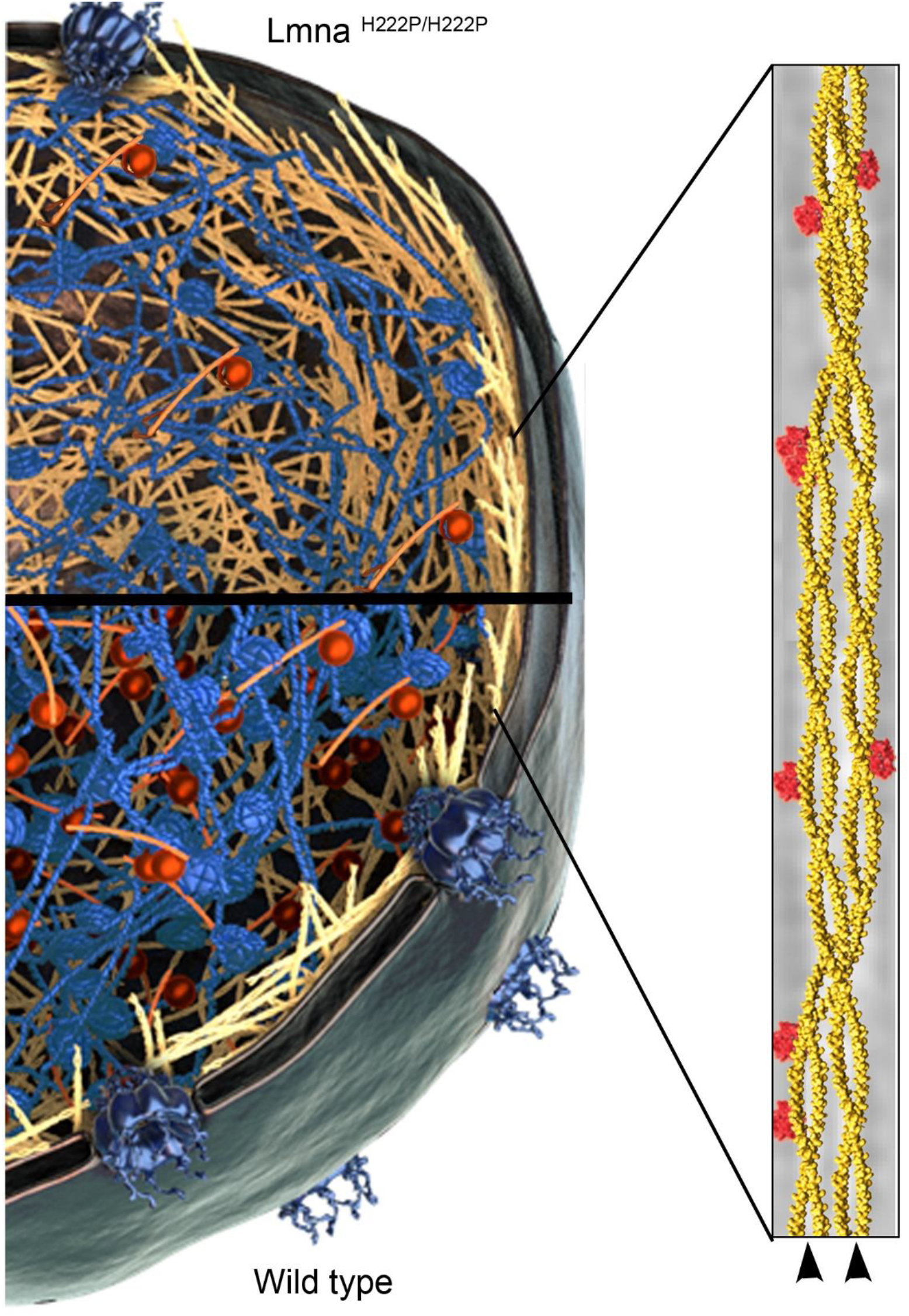
Nuclear envelope and lamins in wild type and *Lmna*_H222P/H222P_ MEFs. The model shows the nuclear lamina (yellow) of wild type MEFs, which has tight interactions with the chromatin (blue) and chromatin binding factors (red). In *Lmna*^H222P/H222P^ MEFs, the nuclear lamin filaments (yellow and tan) are more exposed. The structure of the lamin filaments composed of two head-to-tail lamin dimeric polymers (arrowheads) interacting to form a variety of structures is shown at the right. The model was superimposed on top of reconstituted filaments (Fig. 3); the Ig-like fold domains (reddish) were added subjectively to resemble their appearance in Fig. 3A.

Our results suggest that the distribution of lamin filaments within the lamina of the *Lmna*^H222P/H222P^ MEFs is presumably unaffected. The lamina of most somatic cells is composed of both A- and B-type lamins, although in *Lmna*^H222P/H222P^ fibroblasts, we found that B-type lamin expression was upregulated relative to wild type fibroblasts. We analyzed a large number of lamin filaments, >950,000 filament segments form 233 tomograms, ensuring with a high statistical confidence the close structural resemblance of lamin A/C H222P filaments to the wild type ones. The H222P amino acid substitution in lamin A/C is situated in a linker domain, between coil 1 and 2; therefore, this region presumably can accommodate the mutation without a major structural alteration.

The assembly of lamin dimers into polymers is central for understanding the basic characteristics of the filaments. A recent publication suggested that lamins interact laterally through coil 1 and 2 to form a tetrameric structure (Ahn et al., 2019). This is similar to cytoplasmic IFs (Herrmann and Aebi, 2004). Our experiments revealed the existence of two protofilaments ~1.8 nm in thickness, which have the dimensions of a dimeric coiled-coil structure (Zaccai et al., 2011). This observation supports the notion that two head-to-tail filaments are the basic lamin components that interact to form the mature filaments (Stuurman et al., 1998). Moreover, our *in silico* reconstituted filaments provide evidence that the interactions between the two head-to-tail polymers of dimers are not structurally homogeneous (Fig. 5 arrowheads), allowing lamin filaments to adopt a spectrum of conformations that presumably crucial to their flexibility and the required changes upon the introduction of external forces (Cho et al., 2019; Maurer and Lammerding, 2019; Sapra et al., 2019).

Pathogenic *LMNA* mutations leading to many different amino acid substitutions along the lamin A/C proteins cause striated muscle disease (Briand et al., 2018; Cattin et al., 2013; Maggi et al., 2016; Perrot et al., 2009; Tatli and Medalia, 2018). These mutations appear to result in a loss of some aspect of lamin A/C function, as human patients with *LMNA* mutations leading to haploinsufficiency and *Lmna* null mice both develop striated muscle disease (Bonne et al., 1999; Sullivan et al., 1999). However, how these lamin A/C amino acid substitutions cause cardiomyopathy and muscular dystrophy is poorly understood. Among several hypotheses are that the pathogenic lamin A/C variants lead to altered chromatin organization, gene expression and cellular mechanotransduction (Osmanagic-Myers and Foisner, 2019; Schreiber and Kennedy, 2013). Our results provide structural insights into how a striated muscle disease-causing lamin A/C variant affects nuclear structure. In *Lmna*^H222P/H222P^ MEFs, the overall organization of lamins in the nuclear lamina appears unaffected by the amino acid substitution. However, nuclei are slightly larger with alterations at the lamina-chromatin interface. How these alterations affect the mechanical properties of the nucleus, signaling cascades or gene expression remains to be determined. The normal lamina structure at the resolution of cryo-ET also cannot explain the alterations in nuclear structure observed at the light microscope level in fibroblasts of human subjects with striated muscle disease-causing *LMNA* mutations and transfected cultured cells expressing the pathogenic lamin A variants (Muchir et al., 2004; Ostlund et al., 2001; Raharjo et al., 2001).

Using cryo-ET to examine additional cells expressing different lamin A/C variants could lead to a better understanding of the structure of nuclear lamins and filament assembly. Combining these studies with *in vitro* assembly assays of lamin filaments will lead to a higher-resolution structural resolution of lamins and lamina formation. Examination of cells from mice and human subjects with other mutations causing striated muscle disease and other laminopathies, such as partial lipodystrophy, peripheral neuropathy or progeria, could also provide further insights into pathogenic mechanisms. Furthermore, muscular dystrophy and cardiomyopathy caused by *LMNA* mutations in humans is autosomal dominant. Therefore, the lamin A/C filaments or dimers in the cells of human patients could be composed of both wild type and variant proteins. The autosomal-dominant nature of most human laminopathies would therefore provide an additional challenge in scrutinizing the effects of the disease-causing variants on lamina structure and pathogenic mechanisms.

## Materials and methods

### Preparation and immortalization of MEFs

MEFs were isolated from E14-E15 embryos generated by crosses between *Lmna*^H222P/+^ male and female mice. Briefly, each embryo was cut into fine pieces and incubated at 37°C for 15 min in 1 ml 0.25% trypsin (Gibco, 25200-056). The tissue pieces were then sheared in an 18 gauge needle attached to a syringe and trypsin was inactivated by addition of DMEM supplemented with high glucose, sodium pyruvate, GlutaMAX, and phenol red (Gibco, 10569-010) and containing 15% (v/v) fetal bovine serum (Gibco, 26140-079). Cells were then plated in 10 cm culture dishes and allowed to adhere for 24 h. Non-adherent cells were discarded and the adherent fraction were the MEFs. To establish the immortalized lines, MEFs at passage 2 were infected with a packaged retrovirus, which expresses SV40 large T antigen (a gift from Drs. Eros Lazzerini Denchi and Larry Gerace). Stable immortalized MEF pools were established by selecting the infected cells in 400 μg/ml geneticin (Gibco, 10131035). Genotypes of the embryos and MEFs were confirmed by PCR as previously described (Arimura et al., 2005). The Institutional Animal Care and Use Committee at Columbia University Irving Medical Center approved the protocol.

### Cell culture

Immortalized wild type and *Lmna*^H222P/H222P^ MEFs were cultured in DMEM (Sigma-Aldrich, D5671) supplemented with 10% (v/v) FCS (Sigma-Aldrich, F7524), 1% penicillin-streptomycin (Sigma-Aldrich, P0781), 2 mM l-glutamine (Sigma-Aldrich, G7513), and 400 ug/ml geneticin at 37 °C and 5% CO_2_ in a humidified incubator. Confluent cells were trypsinized (Sigma-Aldrich, T4174) and seeded onto a glow discharged holey carbon covered EM-grids (R 2/1, Au 200, Quantifoil). After approximately 16 h, the grids were subject to the nuclease treatment or they were directly plunged frozen into liquid nitrogen cooled ethane for FIB-milling.

### Knockdown of vimentin in MEFs

Transfection using the committal vimentin shRNAi lentiviral vector (Sigma-Aldrich) and the PolyPlus protocol (Jetprime) yielded 60-80% vimentin knockdown. At 24 h post-transfection, the medium was replaced with DMEM supplement with high glucose, 2 mM l-glutamine, 1% penicillin-streptomycin, 15% (v/v) FCS, sodium pyruvate, 400 μg/ml geneticin and 4 μg/ml puromycin. All untransfected cells died upon the addition of the medium. The transfected cells were expanded and analysed by immunoblotting.

### Quantitative immunofluorescence microscopy

MEFs were seeded onto glass cover slips coated with fibrinogen (50 μg/ml, Sigma-Aldrich, 341576) for 3.5 h in 10% (v/v) FCS. Cells were synchronized by serum starvation for 16 h in 1% FCS. Cells were then fixed in 4% (w/v) paraformaldehyde (Sigma-Aldrich, 16005) for 10 min before being permeabilized in 0.1% Triton X-100 (Sigma-Aldrich, T8787) for 10 min. Permeabilized cells were incubated in a blocking buffer (2 % (w/v) BSA, 22.5 mg/ml glycine in PBS with 0.1 % Tween-20 (PBST)) for 1 h at 25 °C. The cells were incubated in the respective primary antibodies, diluted in blocking buffer, for 1 h at 25 °C. The following primary antibodies were used: lamin A/C (Santa Cruz, sc-376248, 1:100), lamin B1 (Santa Cruz, sc-6217, 1:200), H3K9me3 (Abcam, ab8898, 1:700), H3K27ac (Abcam, ab4729, 1:350). After a 3x 5min wash in PBST, the cells were incubated with the secondary antibodies, diluted in blocking buffer for 1 h at 25 °C. The following secondary antibodies were used: Alexa Fluor 488 donkey anti-goat (Jackson Immuno Research 705-545-003, 1:400), Cy3 donkey anti-rabbit (Jackson Immuno Research 711-165-152, 1:400), Cy3 donkey anti-mouse (Jackson Immuno Research 715-165-150, 1:400). Immunostained cells were washed 3x for 5 min in PBST and 3x for 5 min in PBS before being incubated with Hoechst 33342 (Sigma-Aldrich, B2261) for 15 min at 25 °C. After a final wash 3x for 5 min in PBS, the cover slips were mounted on glass slides with Dako mounting medium (Agilent, S3023), and sealed with nail polish.

Cells were analysed with an automated inverted microscope (Leica Microsystems, DMI4000 B) equipped with a fluorescence lamp and a (Leica Microsystems, DFC365 FX) monochromatic digital camera. Images for quantitative analysis were acquired with a 20x air objective (Leica Microsystems, HCX PL Floutar 20x/0.4). Image analysis was done in ImageJ (Schneider et al., 2012). Threshold of the image was applied manually in the Hoechst channel. Next, the image was used to define the nuclei, their area, and the average signal intensity of each channel obtained with the “Analyse Particles” function. The measured signal intensities for each channel was normalized to the median of the corresponding wild-type and the measured area of the nucleus. For TIRF microscopy, the slides were prepared the same way as described above. TIRF microscopy was performed on an inverted widefield microscope (Leica Microsystems, DMI6000B) equipped with an Andor iXon Ultra 897 EMCCD camera. TIRF images were acquired with an 160x oil objective (Leica Microsystems, HC PL APO 160x/1.43 OIL) and with a penetration depth of 90 nm.

### Immunoblotting

For immunoblotting, MEFs were lysed in 1x cell lysis buffer (Cell Signalling Technology, 9803) with proteinase inhibitor cocktail (Sigma-Aldrich, P8340) and 1mM PMSF (Sigma-Aldrich, 93482-50ML-F). Proteins in the cell lysates were separated by SDS-PAGE. Immunoblots were performed by using anti-lamin B1 (Cance et al., 1992) 1:1,000), anti-lamin B2 (Invitrogen, 33-2100, 1:500), anti-H3K9me3 (Abcam, ab8898, 1:1,000), anti-Lamin A/C (Santa Cruz, SC-20681, 1:5,000), anti-ɣ-tubulin(Sigma-Aldrich, T-5326, 1:1,000), and anti-Gapdh (Ambion, AM4300, 1:5,000) antibodies. Quantification of blots was performed with ImageJ, normalized to loading controls as indicated, and presented as fold change over untreated or wild type MEFs.

### Preparation of ghost nuclei on EM grids

Cells growing on EM-grids were rinsed in PBS supplemented with 2 mM MgCl_2_, permeabilized for 15-20 s in a permeabilization buffer (1X PBS, 0.1% Triton X-100, 600 mM KCl, 10 mM MgCl_2_ and protease inhibitors), and rinsed again. Thereafter, grids were treated with 2.5 units/μl benzonase (Merck, 71206-3) in PBS/2mM MgCl2 for 30 min, washed again prior to applying 3 μl of fiducial gold markers (Aurion, 10 nm, BSA-coated), and then plunged frozen into liquid ethane.

### Cryo-FIB-scanning electron microscopy (SEM) milling

Prior to FIB-milling, the grids were coated with 5 nm Pt/C by using a Leica BAF060 system cooled to −160° C. The grids were transferred to a Zeiss Auriga 40 Crossbeam FIB-SEM. Using the gas injection system, an organometallic platinum protective layer was applied to the grids. Cells were milled with a focused gallium ion beam at a stage temperature of <-150 °C and a stage angle of 18°. The milling was controlled by the NanoPatterning and Visualization Engine software (Zeiss) and observed by SEM. Final thickness of FIB-milled lamellas were 100-200 nm.

### Cryo-ET acquisition

Tilt series were acquired of FIB-milled lamellas and ghost nuclei using an FEI Titan Krios at 300 KeV electron microscope equipped with a Gatan Quantum Energy Filter and a K2 Summit direct electron detection camera. All tilt series were acquired at a magnification of 64,000x and 4-6 μm underfocus with SerialEM (Mastronarde, 2005), resulting in a 0.22 nm/pixel at the specimen level. The data covered an angular range of −60° to 60°, acquired from −30° to +60° followed by −30° to −60°, every 2°, with a total dose of 100 - 140 e^−^/Å^2^. Tilt series acquisition for FIB-milled lamellas were conducted by a dose symmetric tilt scheme (Hagen et al., 2017) from −60° to 60° every 3° with a total dose of 150e^−^/Å^2^.

### Cryo-ET image processing

All tilt series were reconstructed by using the IMOD workflow. For visualization purposes, the tomographic slices picked for the visualization were reconstructed with SIRT algorithm and 8 z slices were projected in slicer window in IMOD. A dataset of 233 tomograms with −4 μm defocus and 120 e^−^/A^2^ were selected for image averaging. The tomograms were reconstructed again in MATLAB using TOM Toolbox. Thereafter, APT workflow (Martins et al., 2020) was followed. Ten manually segmented tomograms (Amira-Avizo 2019.1, Thermo Fischer Scientific) were used as a reference for the neural network segmentation of 233 tomograms. Evenly spaced coordinates every 5 nm along the filaments were picked to determine the centre of each segment along the filament axis. Approximately 967,000 sub-volumes (12×12×4.4 nm^3^) were reconstructed around the picked coordinates and projected to 2D images. The projected images were imported into Relion 3.0. After several rounds of classifications, 200 class averages with a total number of 350,000 particles were obtained. The final 2D class averages were mapped-back two or more sequential segments along a filament and were selected and mapped back to their x-y positions. Surface rendering images of wild type and *Lmna*^H222P/H222P^ MEFs tomograms were generated using the Amira-Avizo 2019.1 software package (Thermo Fischer Scientific).

## Acknowledgements

This work was funded by grants from the Swiss National Science Foundation Grant (SNSF 31003A_179418) and the Mäxi Foundation to O.H. and a grant from the U.S. National Institutes of Health (R01AR04897) to H.J.W. We thank the Center for Microscopy and Image Analysis at the University of Zurich (ZMB), Dr. Yagmur Turgay for technical help in the initial stages and Drs. Eros Lazzerini Denchi and Larry Gerace for reagents and advice on immortalizing MEFs. The authors declare no competing financial interests.

## Authors contributions

H.J.W. and O.M. conceived the project and provide the funding. M.T. and R.T. conducted the experiments and together with M.E. analysed the data. G.B. generated the *Lmna* H222P mice. W.W. bred and cared for mice, generated immortalized MEFs and performed immunoblotting experiments. J-Y. S. generated immortalized MEFs. O.M and H.J.W wrote the manuscript with contributions from all authors.

## Supplementary Materials

**Fig. S1.**
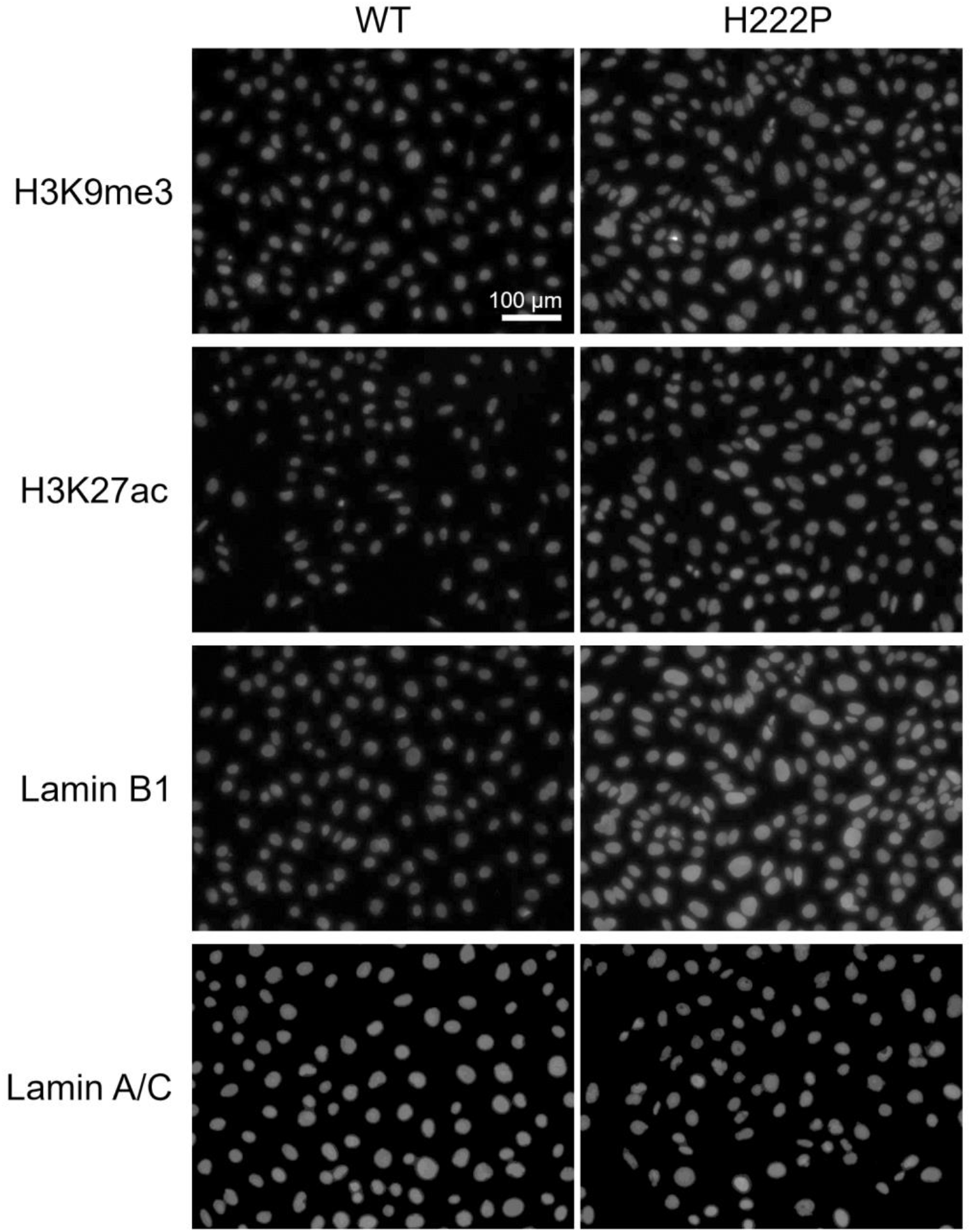
Typical immunofluorescence images used for quantification of H3K9me3, H3K27ac, lamin B1 and lamin A/C shown in Figure 1. Immunofluorescent images were acquired and analysed as described in Materials and methods.

**Fig. S2.**
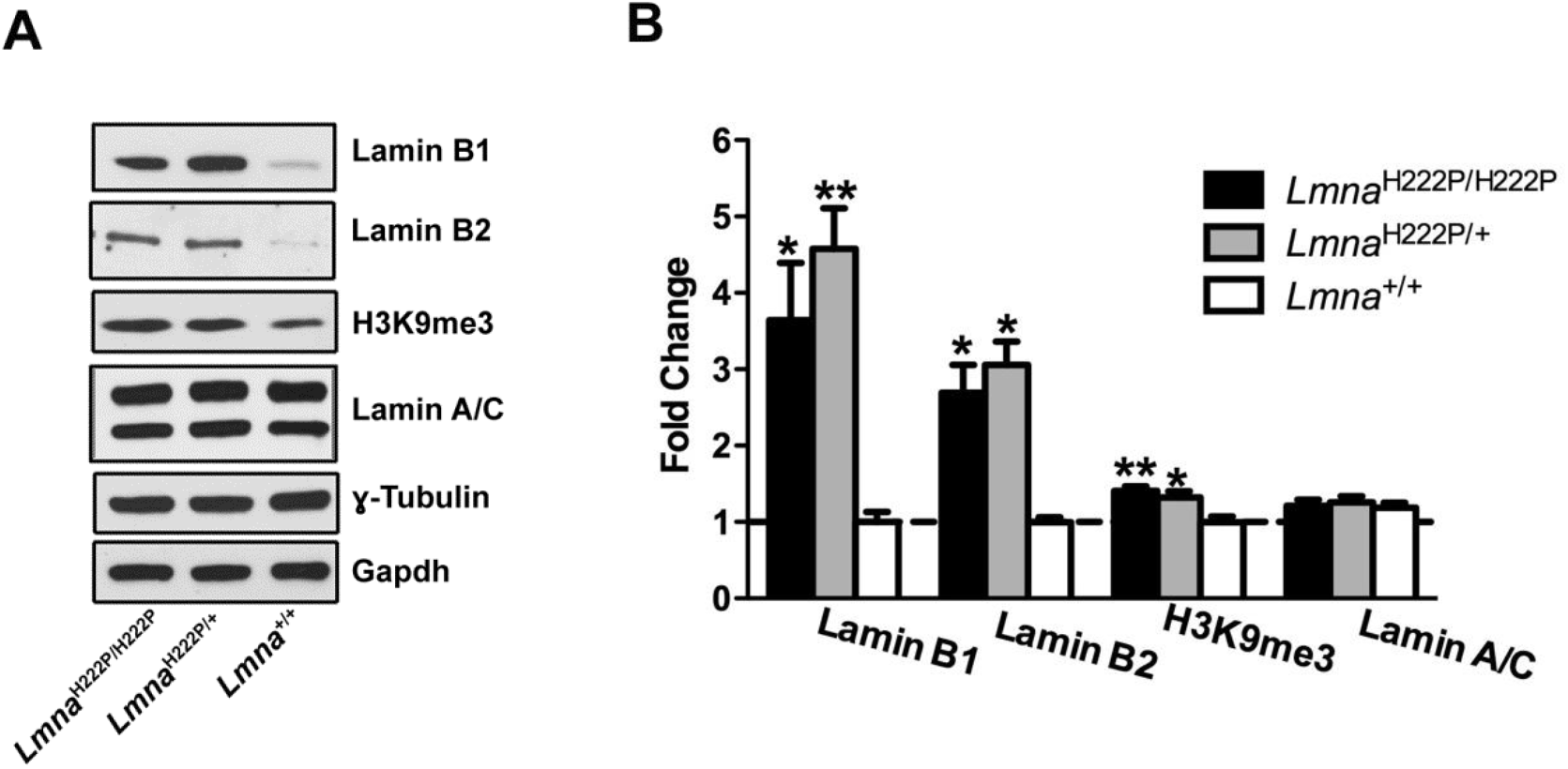
Endogenous protein expression levels in immortalized MEFs. **A**. Representative immunoblot of endogenous lamin B1, lamin B2, histone H3K9me3 and lamin A/C expression in immortalized *Lmna*^H222P/H222P^, *Lmna*^H222P/+^ and wild type (*Lmna*^+/+^) MEFs. ɣ-Tubulin and Gapdh are loading controls. **B.** Quantification of lamin B1, lamin B2, histone H3K9me3 and lamin A/C normalized to ɣ-tubulin and Gapdh presented as fold-change over wild type, n = 3. *p<0.05 and **p<0.005 compared to wild type by one way ANOVA Tukey’s multiple comparison test.

**Fig. S3.**
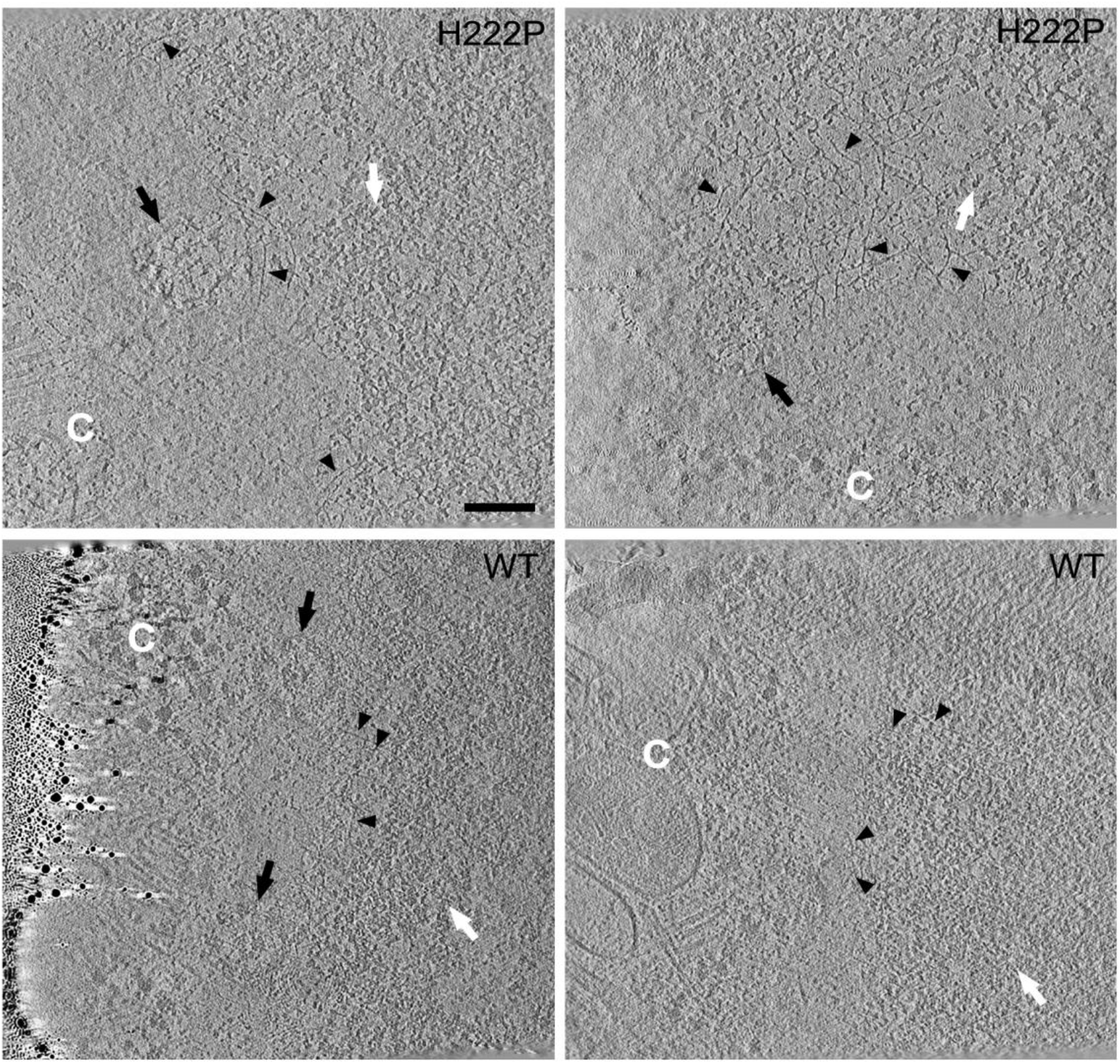
Cryo-ET slices through wild type and *Lmna*_H222P/H222P_ MEFs. Cryo-ET was applied to cryo-FIB milled lamellea of *Lmna*^H222P/H222P^ MEFs (H222P) and wild type (WT) MEFs. X-Y slices, 7 nm in thickness, through the nuclear envelopes of four nuclei are shown. The nuclear pore complexes (black arrows) mark the position of the nuclear membranes. Lamin filaments (black arrowheads) are detected in between the cytoplasm (C) and nucleosomes (white arrows). In the wild type cells, the chromatin densities approach the lamin filaments, partly blocking their visualization. Scale bar 100 nm

**Fig. S4.**
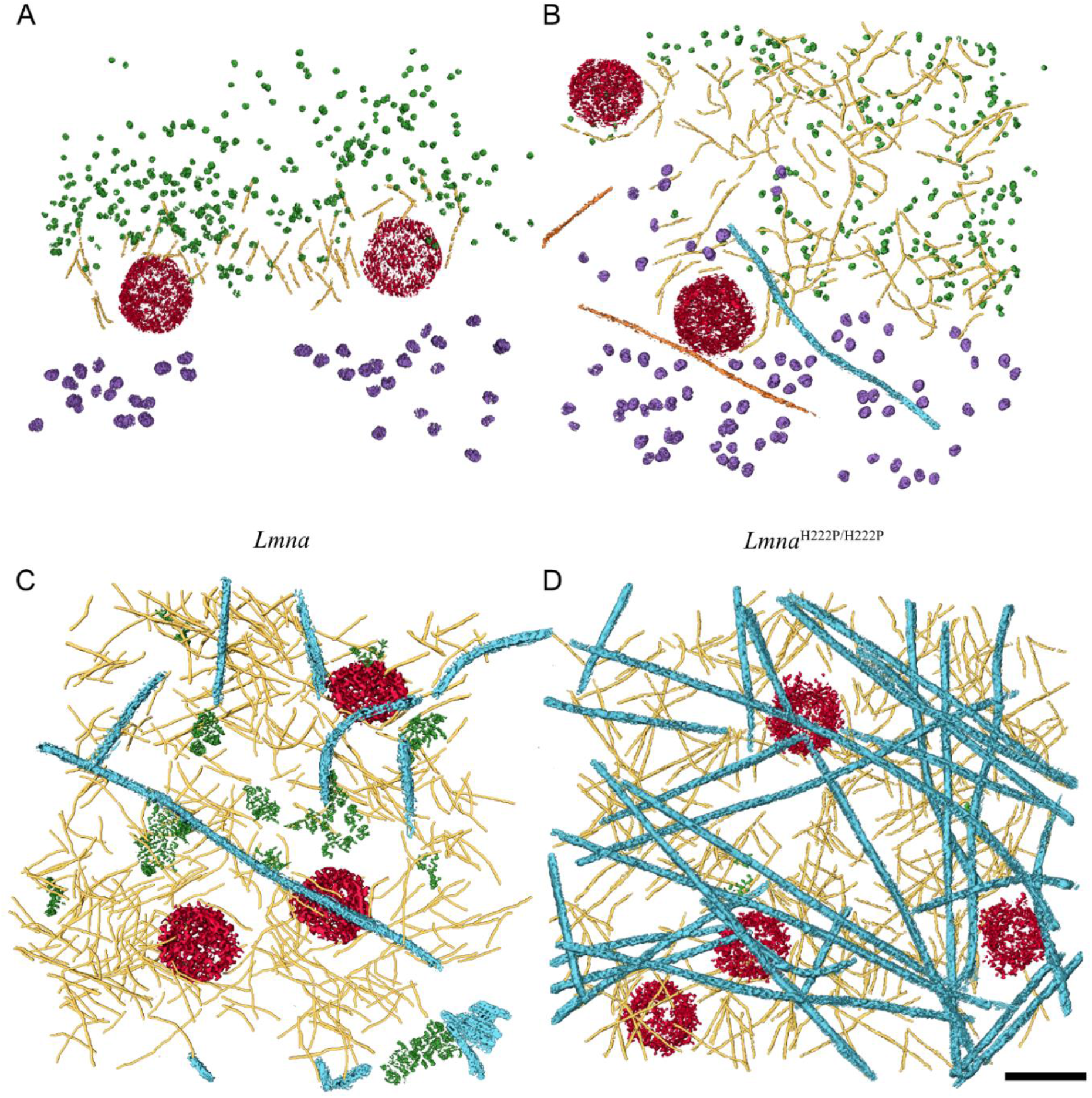
Rendered views of tomograms of FIB-SEM lamellae and ghost nuclei from wild type and *Lmna*_H222P/H222P_ MEFs. **A.** Rendered view of cryo-tomogram of cryo-FIB milled lamellae from wild type MEFs. **B.** Rendered view of cryo-tomogram of cryo-FIB milled lamellae from *Lmna*^H222P/H222P^ MEFs. The high density of the nucleosomes (green) at the nuclear envelope of the wildtype cells, the identification of lamins is less efficient. The lamellae exhibit a thickness of 150nm. **C.** Rendered view of cryo-tomogram from ghost nucleus from wild type MEFs (WT). **D.** Rendered view of cryo-tomogram from ghost nucleus from *Lmna*^H222P/H222P^ MEFs (H222P). Lamin filaments (yellow), cytoplasmic vimentin filaments (turquoise), nuclear pore complexes (red) and nucleosomes (green) are shown, while ribosomes (purple), actin filaments (orange) were frequently observed in the in situ measurements (FIB-SEM milling approach). Because of the lower density of nuclear structures at the nuclear envelope of the lamin A/C H222P mutated nuclei, the density of lamin filaments can be detected and followed resulted in a better resolved structures. Scale bar 100 nm

**Fig. S5.**
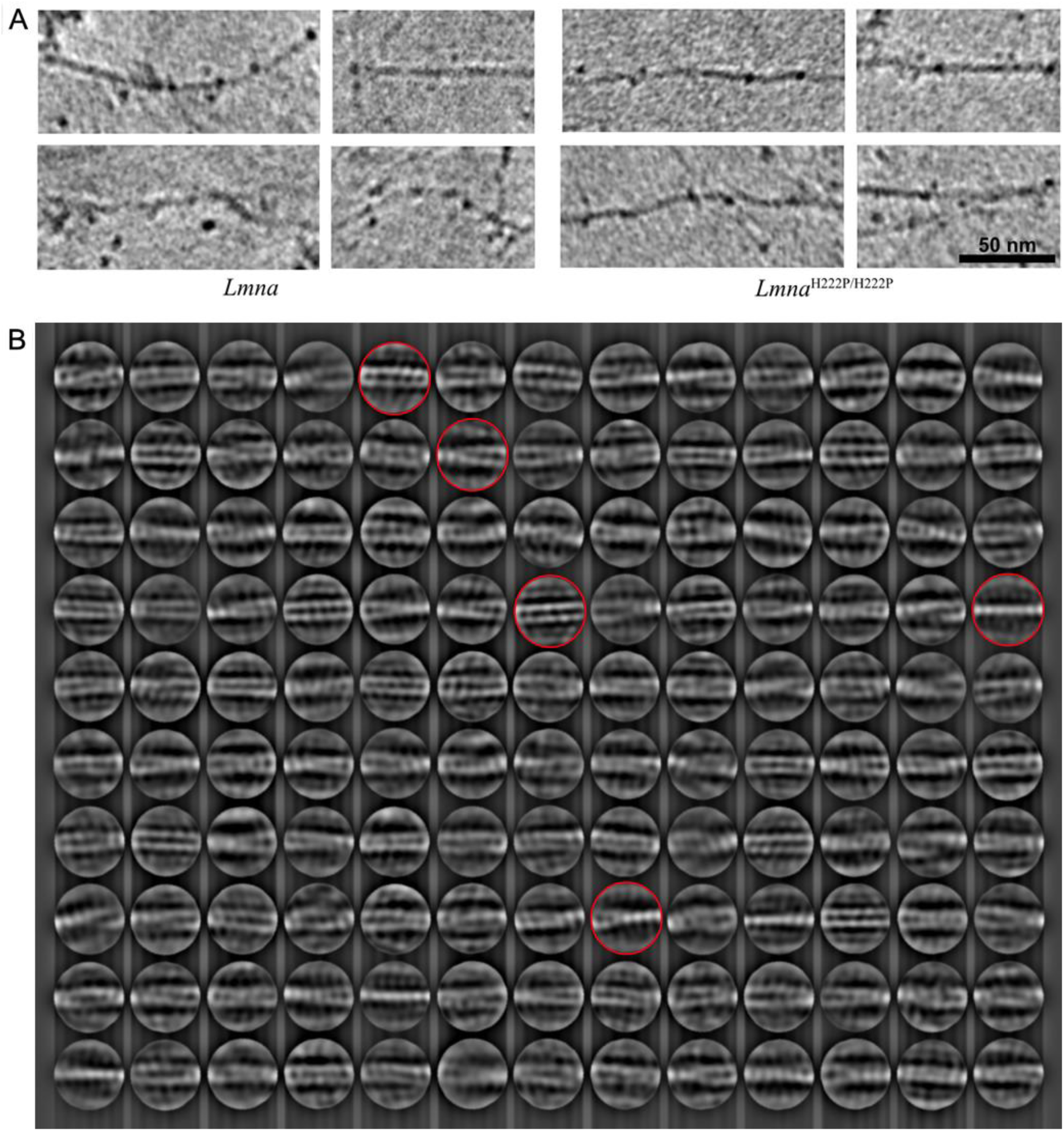
Analysis of lamin filaments in ghost nuclei. **A.** Lamin filaments found in cryo-tomograms of ghost nuclei and from wild type and *Lmna*^H222P/H222P^ MEFs. The filaments are similar in their appearance although less material was detected around the filaments in *Lmna*^H222P/H222P^ MEF ghost nuclei. **B.** Two-dimensional class averages were obtained and displayed. The classes shown in Fig. 4B are indicated by red circles.

